# Space-time clusters of dengue, chikungunya, and Zika cases in the city of Rio de Janeiro

**DOI:** 10.1101/521591

**Authors:** Laís Picinini Freitas, Oswaldo Gonçalves Cruz, Rachel Lowe, Marilia Sá Carvalho

## Abstract

Brazil is a dengue-endemic country where all four dengue virus serotypes circulate and cause seasonal epidemics. Recently, chikungunya and Zika viruses were also introduced. In Rio de Janeiro city, the three diseases co-circulated for the first time in 2015-2016, resulting in what is known as the ‘triple epidemic’. In this study, we identify space-time clusters of dengue, chikungunya, and Zika, to understand the dynamics and interaction between these simultaneously circulating arboviruses in a densely populated and heterogeneous city.

We conducted a spatio-temporal analysis of weekly notified cases of the three diseases in Rio de Janeiro city (July 2015 – January 2017), georeferenced by 160 neighbourhoods, using Kulldorff’s scan statistic with discrete Poisson probability models.

There were 26549, 13662, and 35905 notified cases of dengue, chikungunya, and Zika, respectively. The 17 dengue clusters and 15 Zika clusters were spread all over the city, while the 14 chikungunya clusters were more concentrated in the North and Downtown areas. Zika clusters persisted over a longer period of time. The multivariate scan statistic – used to analyse the three diseases simultaneously – detected 17 clusters, nine of which included all three diseases.

This is the first study exploring space-time clustering of dengue, chikungunya, and Zika in an intraurban area. In general, the clusters did not coincide in time and space. This is probably the result of the competition between viruses for host resources, and of vector-control attitudes promoted by previous arbovirus outbreaks. The main affected area – the North region – is characterised by a combination of high population density and low human development index, highlighting the importance of targeting interventions in this area. Spatio-temporal scan statistics have the potential to direct interventions to high-risk locations in a timely manner and should be considered as part of the municipal surveillance routine as a tool to optimize prevention strategies.

**Author summary:** Dengue, an arboviral disease transmitted by *Aedes* mosquitoes, has been endemic in Brazil for decades, but vector-control strategies have not led to a significant reduction in the disease burden and were not sufficient to prevent chikungunya and Zika entry and establishment in the country. In Rio de Janeiro city, the first Zika and chikungunya epidemics were detected between 2015-2016, coinciding with a dengue epidemic. Understanding the behaviour of these diseases in a triple epidemic scenario is a necessary step for devising better interventions for prevention and outbreak response. We applied scan statistics analysis to detect spatio-temporal clustering for each disease separately and for all three simultaneously. In general, clusters were not detected in the same locations and time periods, possibly due to competition between viruses for host resources, and change in behaviour of the human population (e.g. intensified vector-control activities in response to increasing cases of a particular arbovirus). Neighbourhoods with high population density and social vulnerability should be considered as important targets for interventions. Particularly in the North region, where clusters of the three diseases exist and the first chikungunya cluster occurred. The use of space-time cluster detection can direct intensive interventions to high-risk locations in a timely manner.

## Introduction

Dengue has been endemic in Brazil for more than 30 years. Since 2010, all four dengue virus (DENV) serotypes circulate in the country [1]. The first chikungunya and Zika outbreaks in Brazil were detected in 2014 and 2015, respectively, both in the Northeast region. In 2016, 1.5 million dengue cases, 270 thousand chikungunya cases, and more than 200 thousand Zika cases were notified in the country [2]. Initially described as a benign disease, Zika quickly became a serious public health problem after the association of the disease during pregnancy with congenital malformations, such as microcephaly, was discovered [3–5].

The co-circulation of DENV, chikungunya virus (CHIKV) and Zika virus (ZIKV), poses a serious public health and economic burden [6,7]. The Brazilian government has implemented dengue prevention and control measures in the form of vector-control interventions, but there is no evidence that vector-control has had a significant effect in reducing transmission in Brazil or other parts of the world [8]. The widespread presence of the vector (mainly *Aedes aegypti* but also *Aedes albopictus*), a highly mobile population, and low or lack of herd immunity resulted in simultaneous and overlapping outbreaks of all three diseases, a phenomenon that has been referred to as the ‘triple epidemic’. Understanding the behaviour of dengue, Zika, and chikungunya, when they compete in time and space, is a step forward in improving the design of interventions for prevention and outbreak response [9].

The Brazilian National Notifiable Diseases Information System (Sistema de Vigilância de Agravos de Notificação [SINAN]) is the Ministry of Health’s system for surveillance of diseases included in the national list of compulsory notification. Dengue has been a notifiable disease since 1961, and chikungunya since 2011. Zika was only included in February 2016, but since June 2015 Zika was monitored through sentinel surveillance [10,11]. Most notifications are made by physicians working in public health facilities, based on diagnostic protocols by the Ministry of Health. SINAN receives a large number of notifications and it thought to accurately represent the overall trend of the dengue situation in Brazil [12].

Considering DENV, CHIKV, and ZIKV share the same vectors and human hosts, we conducted a spatio-temporal analysis of notified cases to identify clusters and understand the dynamics of these diseases in a scenario of triple epidemics. Rio de Janeiro was the chosen city for this analysis for the following reasons: a history of large dengue epidemics with sustained transmission; the recent occurrence of CHIKV and ZIKV epidemics in 2015-2016; co-circulation of DENV, CHIKV and ZIKV; a high number of reported cases; the possibility to work with georeferenced cases in an intra-urban context; multiple environmental settings within the city; high human mobility; vector abundance; and health professionals experienced in dealing with dengue as a result of the epidemiological scenario.

## Methods

### Study site

Rio de Janeiro is the second largest city in Brazil, with approximately 6,3 million inhabitants (2010 census), 1204 km^2^ and 160 neighbourhoods (Fig 1). The city has the 45^th^highest Human Development Index (HDI) of the country, of 0.799 (varying from 0.604 to 0.959 inside the city) [13,14]. The population density is 5249 inhabitants per km^2^. Rio de Janeiro has a tropical climate, with temperature and rainfall varying depending on altitude, vegetation and ocean proximity. The average annual temperature is 23.7°C, and the annual accumulated precipitation is 1069 mm. During the summer months (December to March), high temperatures (around 40ºC) and thunderstorms are common [15].

**Fig 1.**
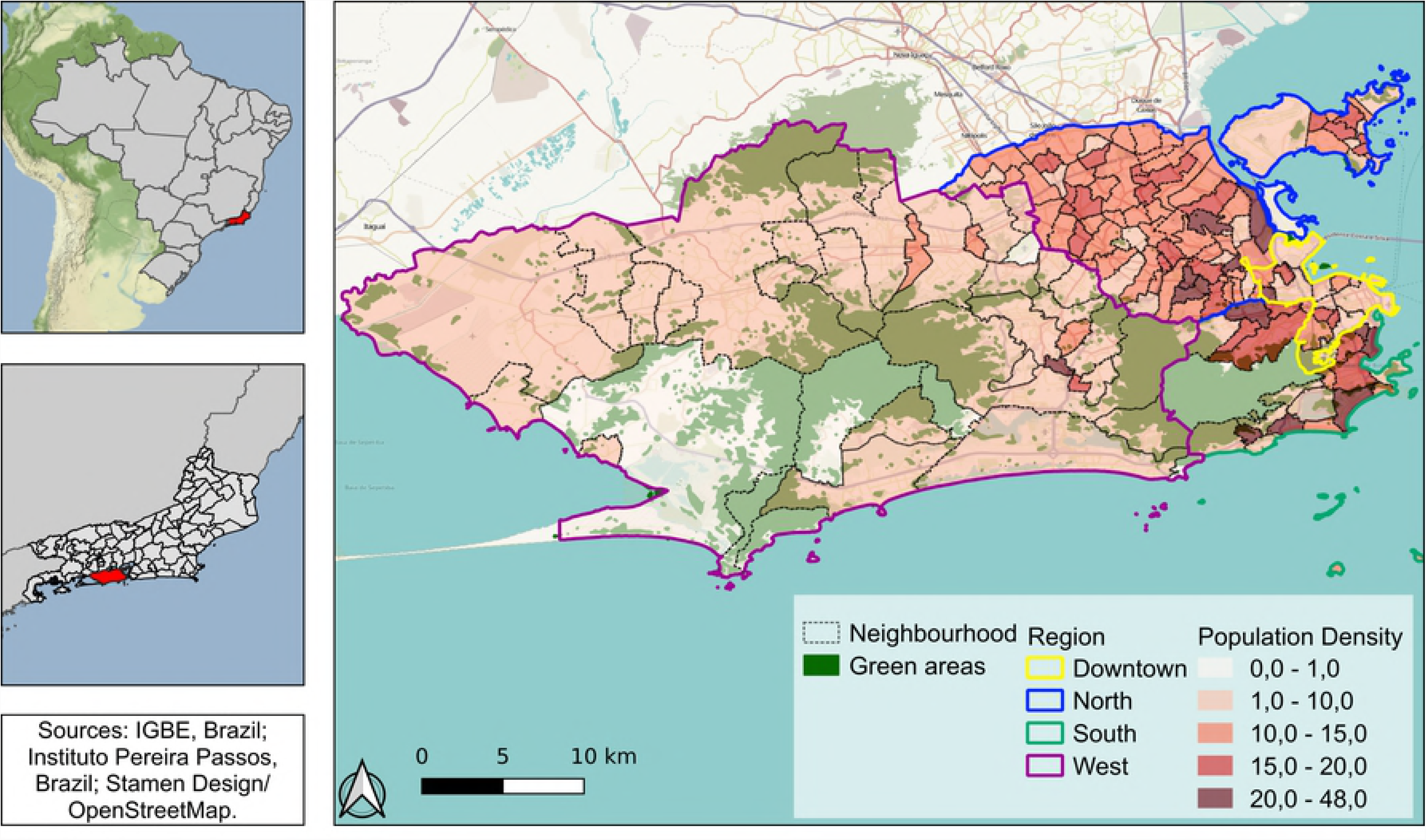
Rio de Janeiro city population density and green areas, by region and neighbourhood, 2010. Map created using QGIS (version 3.4.3). Sources: Brazilian Institute of Geography and Statistics (IBGE) and Instituto Pereira Passos – Rio de Janeiro City Hall, Brazil. Base map from Stamen Design and Open Street Maps.

The 160 neighbourhoods are grouped into four large regions (North, South, Downtown and West), reflecting the geographical position and history of occupation. Almost all neighbourhoods are a mixture of very poor slums (“favelas”) and more affluent areas of residence. The North region is very urbanized, with high population density, few green areas and very large favelas. Nearly 27% of the population of this region, almost 2.4 million people, lived in favelas in the 2010 demographic census [16]. The South region is the most popular tourist destination in Rio de Janeiro, with famous beaches, green areas, and neighbourhoods with the highest HDI of the city [13]. The Downtown region is the historical, commercial and financial center of the city, with many green areas and cultural centers. Finally, the West region has been urbanized and populated more recently, and is less densely populated [15].

### Data

Data on dengue, chikungunya, and Zika cases were obtained from SINAN via the Rio de Janeiro Municipal Secretariat of Heath, and are publicly available. The Municipal Secretariat of Health georeferenced 91% of dengue cases, 95% of chikungunya cases and 92% of Zika cases.

We analysed all cases of dengue, Zika and chikungunya occurring in Rio de Janeiro municipality between 27 July 2015 and 21 January 2017 (epidemiological weeks 30-2015 and 03-2017), grouped by epidemiological week and neighbourhood of residence. Population data by neighbourhood and shapefiles were obtained from the Instituto Pereira Passos (available at: http://www.data.rio/).

### Space-time analysis

For spatio-temporal detection of clusters, Kulldorff’s scan statistic with a discrete Poisson probability model was applied for each disease individually and for the three diseases simultaneously (multivariate scan statistic with multiple data sets). The scan statistic uses moving cylinders across space (i.e. the base of the cylinder) and time (i.e. the height of the cylinder) to identify clusters, by comparing the observed number of cases inside the cylinder to the expected number of cases [17,18]. The detected clusters are ordered in the results section according to the likelihood ratio, such that the cluster with the maximum likelihood ratio is the most likely cluster, that is, the cluster least likely to be due to chance. The relative risk for each cluster is calculated as the observed number of cases within the cluster divided by the expected number of cases within the cluster, divided by the observed number of cases outside the cluster divided by the expected number of cases outside the cluster [19].

The multivariate scan statistic for multiple data sets was applied to simultaneously search for clusters of dengue, Zika and chikungunya that coincided in time and space. This technique calculates for each window the log likelihood ratio for each disease. Then, the likelihood for a particular window is calculated as the sum of the log likelihood ratios for the diseases with more than the expected number of cases. In the same way as for a single disease, the maximum of all the summed log likelihood ratios constitutes the most likely cluster [19,20].

For each model, Monte Carlo simulations (n=999) were performed to assess statistical significance. We considered statistically significant clusters (p-value < 0.05) that did not coincide in space (with no geographical overlap) and that included a maximum of 50% of the population of the city (nearly 3,1 million people). With only these parameters, two large clusters covering most of the city were detected (S1 Fig A), which is not useful if we are interest in identifying risk areas to direct interventions. After testing several combinations of temporal and spatial parameters (such as the size of the temporal window and maximum population at risk inside the cluster), we chose the combination that resulted in a reasonable number of clusters that aggregated close together and in similar locations that could also be targeted for local interventions (S1 Fig). The temporal window was set to be at least 1 week and a maximum of 4 weeks. Clusters were restricted to have at least 5 cases. In the output parameters, clusters were restricted to include a maximum of 5% of the population of the city (nearly 315 thousand people).

SaTScan™ (version 9.5, https://www.satscan.org/) software was applied within R (version 3.4.4, https://www.r-project.org/), using the package rsatscan (version 0.3.9200) [21–23]. Maps were produced using the ggplot2 (version 3.1.0) package in R [24].

## Results

In Rio de Janeiro, between 27 July 2015 and 21 January 2017 (epidemiological weeks 30-2015 and 03-2017), 76116 cases of dengue, chikungunya, and Zika were reported (Table 1). More than 85% of neighbourhoods had at least 10 cases of each disease. Zika presented the highest number of notifications, resulting in an incidence of 568.1 cases per 100000 inhabitants. Most cases occurred between December 2015 and June 2016 (88.5%). The epidemic curves differed slightly in time, with high incidence of all three diseases between April and June 2016 (Fig 2). In March 2016, Zika cases started to decrease while dengue and chikungunya cases were still on the increase. While dengue and Zika were active by the end of 2015, chikungunya cases only started to rise in March 2016. Notifications of the three diseases declined after May. Interestingly, the shape of the Zika epidemic curve does not have a clear peak.

**Table 1.** Notifications of dengue, chikungunya, and Zika cases between epidemiological weeks 30-2015 and 03-2017 in Rio de Janeiro city, Brazil.

|  | Dengue | Chikungunya | Zika |
| --- | --- | --- | --- |
| Total number of cases | 26549 | 13662 | 35905 |
| Incidence per 100000 inhabitants | 420.0 | 216.2 | 568.1 |
| Maximum n° of cases per week | 2094 | 1101 | 1799 |
| Week with maximum n° of cases | 14-2016 | 17-2016 | 07-2016 |
| N° of neighbourhoods with at least 1 case | 158 | 159 | 160 |
| N° of neighbourhoods with at least 10 cases | 147 | 136 | 155 |

**Fig 2.**
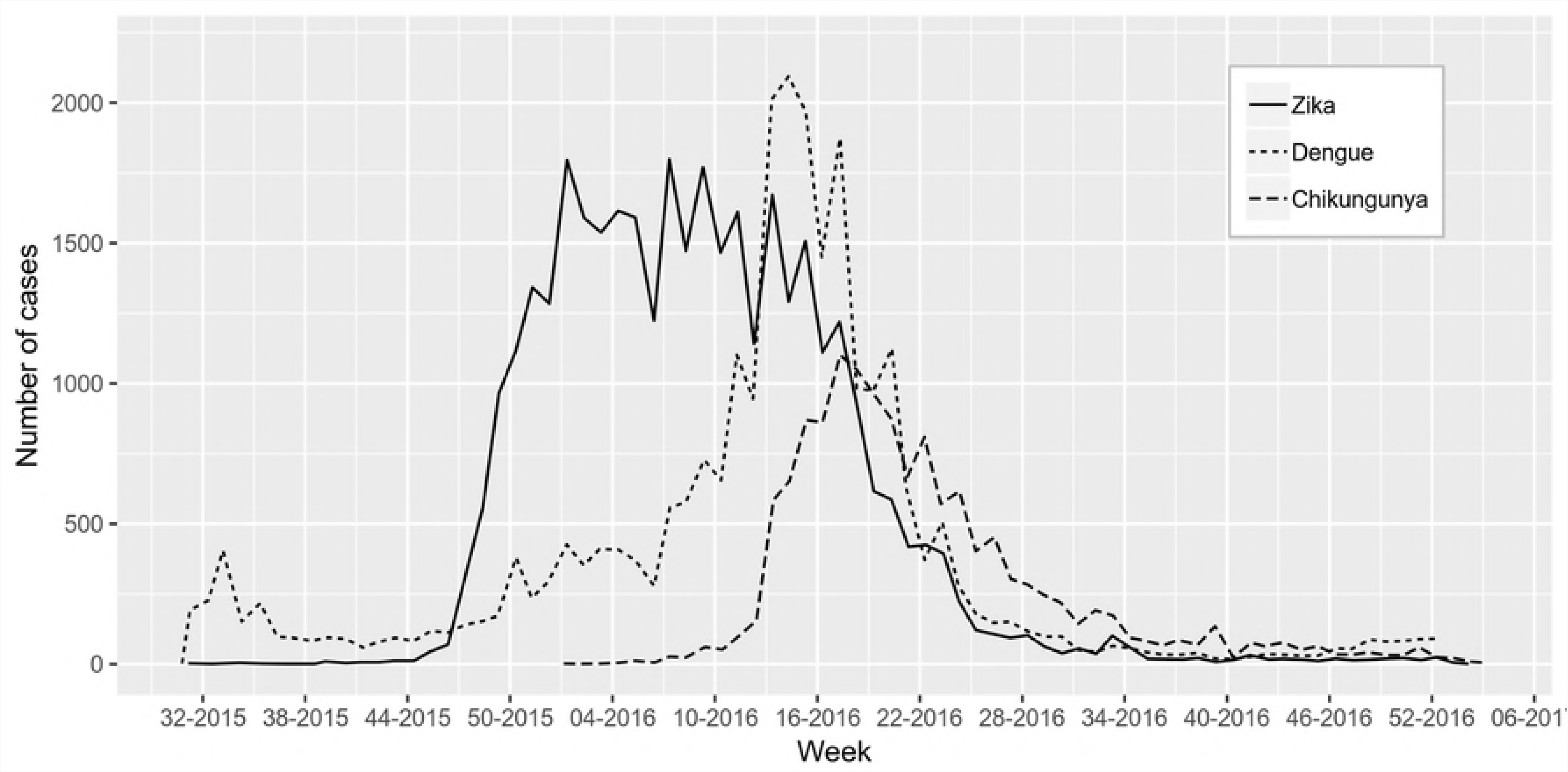
Number of reported dengue (dotted line), chikungunya (dashed line), and Zika (solid line) cases between 27 July 2015 and 21 Jan 2017, Rio de Janeiro city, Brazil. Source: Sistema de Vigilância de Agravos de Notificação (SINAN) – Ministry of Health, Brazil.

### Dengue cases clusters

Scan statistics detected 17 dengue cases clusters (Table 2). Clusters were detected in different parts of the city (Fig 3A). The most likely cluster was located in the North zone of Rio de Janeiro city. Cluster 2 contained only one neighbourhood in the Downtown area with a relative risk of 172.67 (S2 Fig A). Clusters were detected within a short time period, from March to May 2016, except for cluster 15 that started in December 2015 (Fig 3B). The first dengue cluster in time was detected in the West zone (S3 Fig A).

**Table 2.** Characteristics of dengue clusters between epidemiological weeks 30-2015 and 03-2017, Rio de Janeiro city, Brazil. Clusters are ordered according to the maximum likelihood ratio, with 1 being the most likely cluster.

| Cluster | Time period (week) | Observed cases | Population | Relative risk |
| --- | --- | --- | --- | --- |
| 1 | 10 to 14-2016 | 1081 | 293943 | 17.56 |
| 2 | 12 to 16-2016 | 464 | 12556 | 172.67 |
| 3 | 13 to 17-2016 | 905 | 296392 | 14.48 |
| 4 | 13 to 17-2016 | 692 | 243125 | 13.39 |
| 5 | 11 to 15-2016 | 528 | 178123 | 13.87 |
| 6 | 13 to 17-2016 | 425 | 105515 | 18.78 |
| 7 | 13 to 17-2016 | 438 | 296540 | 6.88 |
| 8 | 12 to 16-2016 | 363 | 304235 | 5.54 |
| 9 | 16 to 17-2016 | 170 | 238838 | 13.15 |
| 10 | 13 to 17-2016 | 156 | 94626 | 7.61 |
| 11 | 12 to 16-2016 | 249 | 273908 | 4.20 |
| 12 | 10 to 14-2016 | 184 | 156688 | 5.42 |
| 13 | 14 to 18-2016 | 34 | 3361 | 46.50 |
| 14 | 12 to 15-2016 | 116 | 187930 | 3.79 |
| 15 | 52-2015 to 4-2016 | 79 | 101443 | 3.58 |
| 16 | 13 to 17-2016 | 147 | 311869 | 2.17 |
| 17 | 12 to 14-2016 | 30 | 69356 | 3.98 |

**Fig 3.**
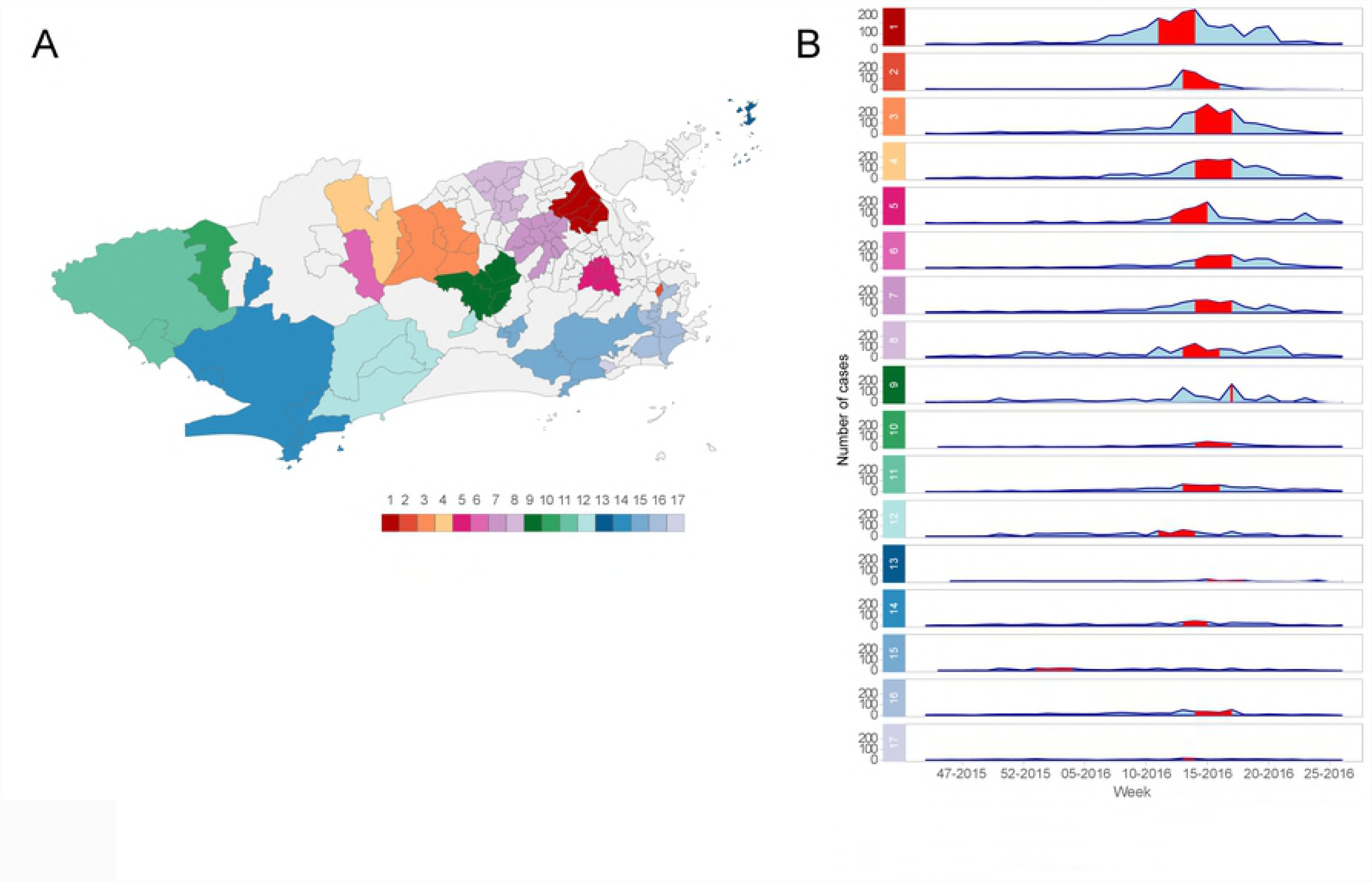
(A) Dengue cases clusters and (B) temporal distribution of dengue cases by cluster, between epidemiological weeks 30-2015 and 03-2017, Rio de Janeiro city, Brazil. Map created using R (version 3.4.4) with ggplot2 package (version 3.1.0). Sources: Sistema de Vigilância de Agravos de Notificação (SINAN) – Ministry of Health, Brazil, and Instituto Pereira Passos – Rio de Janeiro City Hall, Brazil.

### Chikungunya cases clusters

For chikungunya, 14 clusters were detected (Table 3). Unlike dengue, chikungunya clusters were rarely seen in the West of Rio de Janeiro city, with clusters detected in only 7 neighbourhoods of this region (Fig 4A, clusters 6, 9 and 13). The most likely cluster was located in the Downtown of Rio de Janeiro city and had the highest relative risk (S2 Fig B). Clusters were also detected within a restricted time period, between 27 March and 11 June (Fig 4B). The first chikungunya cluster in time occurred in the northern border of the city (S3 Fig B).

**Table 3.**
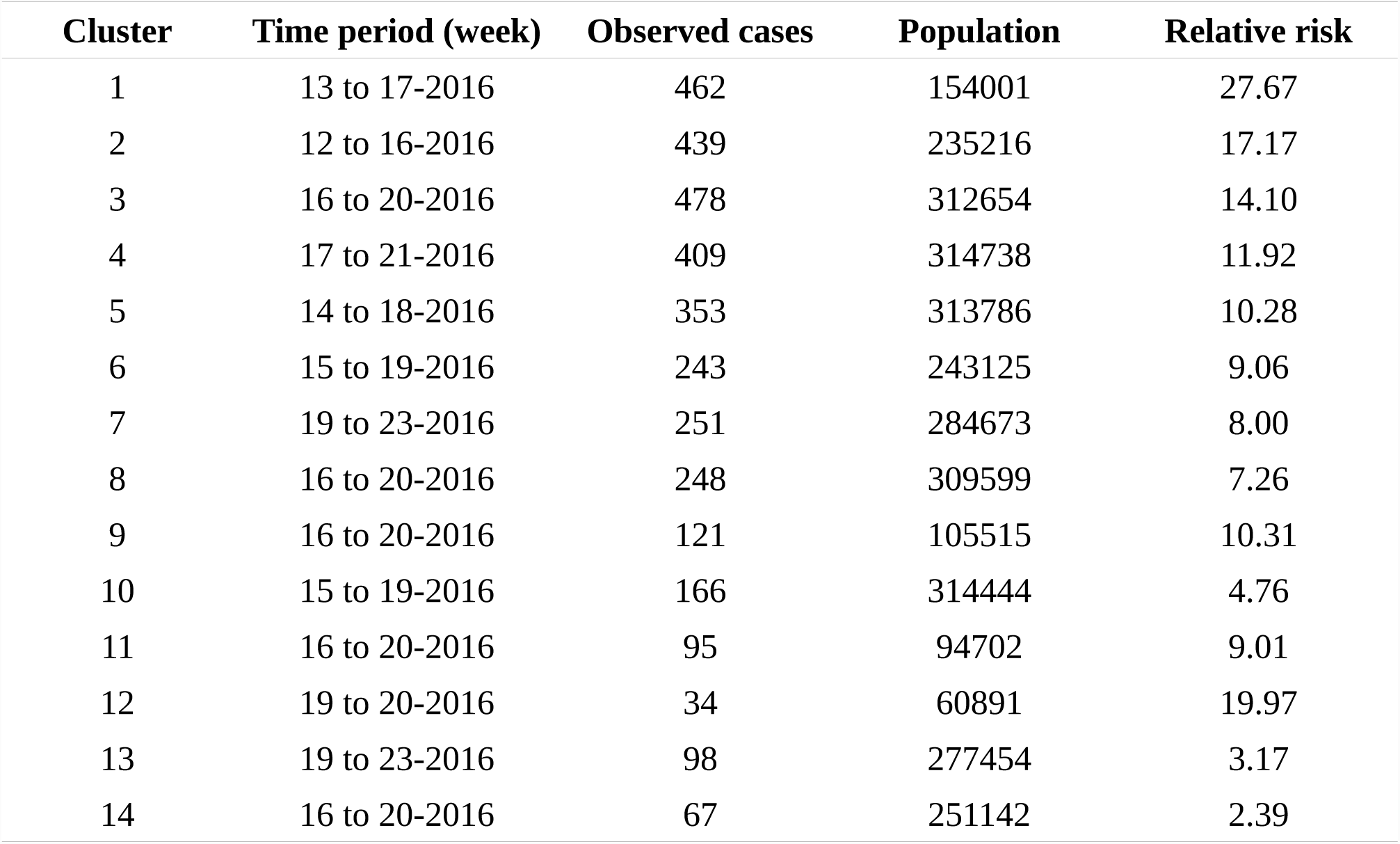
Characteristics of chikungunya clusters between epidemiological weeks 30-2015 and 03-2017, Rio de Janeiro city, Brazil. Clusters are ordered according to the maximum likelihood ratio, with 1 being the most likely cluster.

**Fig 4.**
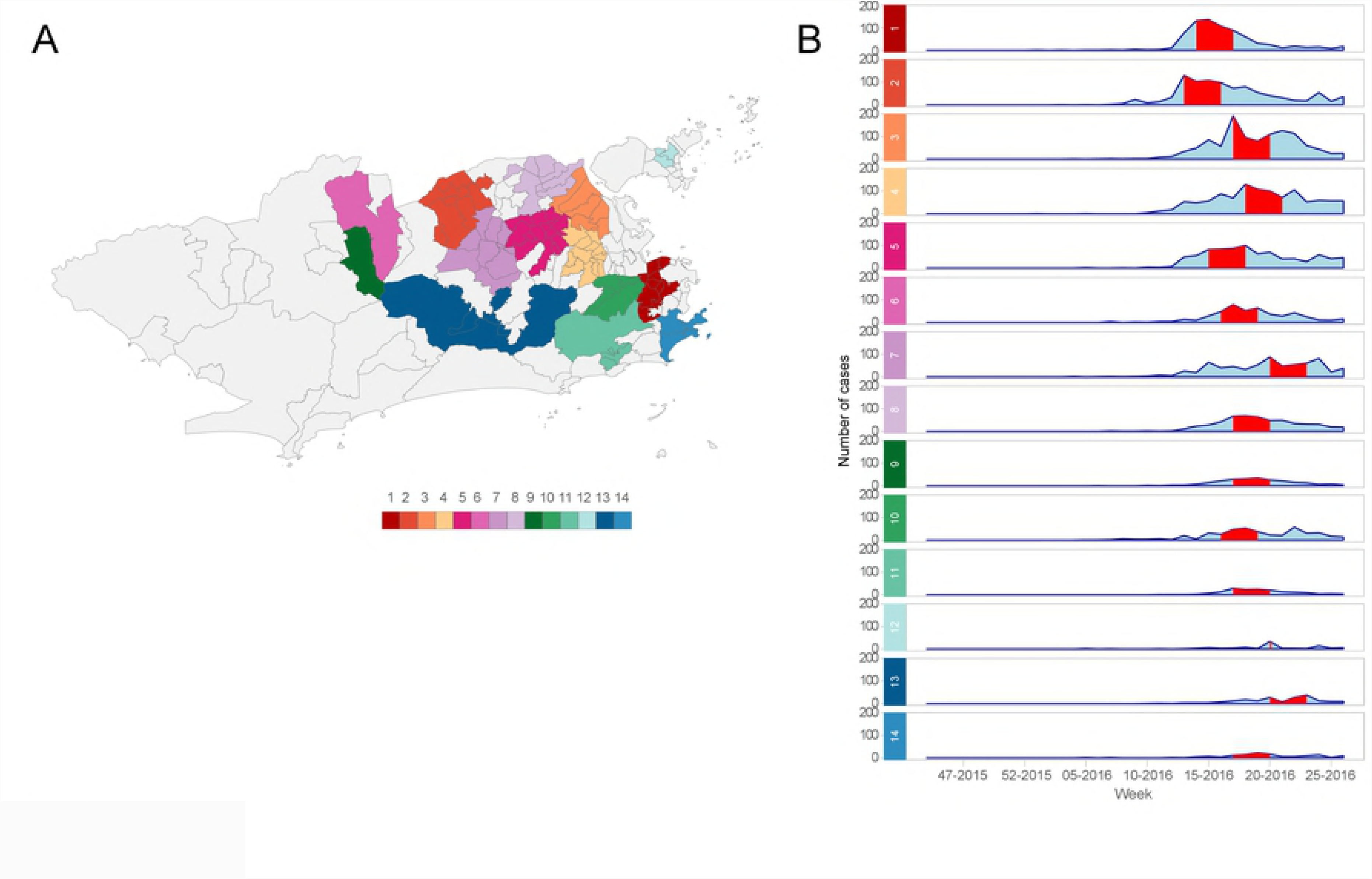
(A) Chikungunya cases clusters and (B) temporal distribution of chikungunya cases by cluster, between epidemiological weeks 30-2015 and 03-2017, Rio de Janeiro city, Brazil. Map created using R (version 3.4.4) with ggplot2 package (version 3.1.0). Sources: Sistema de Vigilância de Agravos de Notificação (SINAN) – Ministry of Health, Brazil, and Instituto Pereira Passos – Rio de Janeiro City Hall, Brazil.

### Zika cases clusters

There were 15 Zika clusters, distributed all over the city, similar to the observed pattern for dengue (Fig 5A, Table 4). The most likely cluster was located in the West of Rio de Janeiro city, a region where chikungunya clusters were rarely observed. This cluster also had the highest relative risk (S2 Fig C). In contrast to dengue and chikungunya, Zika clusters occurred over a longer period of time, between December 2015 and May 2016 (Fig 5B). The third most likely cluster occurred 8 weeks after the first one. The first Zika clusters in time emerged in the North of the city (S3 Fig C).

**Table 4.** Characteristics of Zika clusters between epidemiological weeks 30-2015 and 03-2017, Rio de Janeiro city, Brazil. Clusters are ordered according to the maximum likelihood ratio, with 1 being the most likely cluster.

| Cluster | Time period (week) | Observed cases | Population | Relative risk |
| --- | --- | --- | --- | --- |
| 1 | 52-2015 to 4-2016 | 739 | 179689 | 14.23 |
| 2 | 49-2015 to 1-2016 | 496 | 236282 | 7.21 |
| 3 | 12 to 16-2016 | 517 | 275257 | 6.46 |
| 4 | 1 to 5-2016 | 545 | 309349 | 6.06 |
| 5 | 50-2015 to 1-2016 | 408 | 277724 | 6.71 |
| 6 | 13 to 17-2016 | 480 | 307234 | 5.36 |
| 7 | 6 to 10-2016 | 358 | 170799 | 7.18 |
| 8 | 6 to 10-2016 | 389 | 231774 | 5.75 |
| 9 | 49-2015 to 1-2016 | 426 | 294447 | 4.96 |
| 10 | 15 to 18-2016 | 355 | 297833 | 5.44 |
| 11 | 48 to 52-2015 | 362 | 298052 | 4.16 |
| 12 | 3 to 7-2016 | 314 | 233051 | 4.61 |
| 13 | 6 to 10-2016 | 347 | 289188 | 4.10 |
| 14 | 7 to 11-2016 | 357 | 306508 | 3.98 |
| 15 | 50-2015 to 2-2016 | 112 | 72058 | 5.29 |

**Fig 5.**
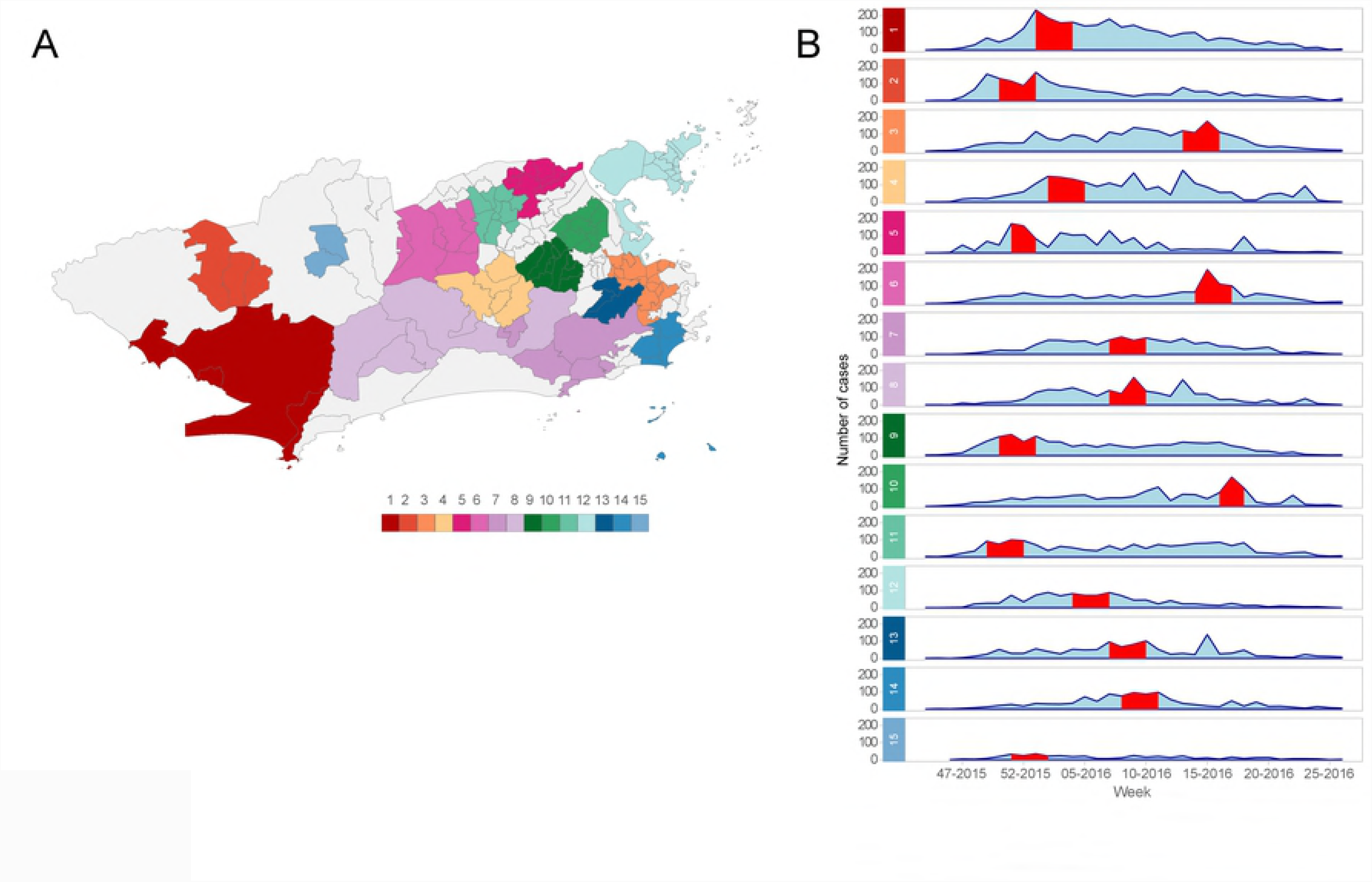
(A) Zika cases clusters and (B) temporal distribution of Zika cases by cluster, between epidemiological weeks 30-2015 and 03-2017, Rio de Janeiro city, Brazil. Map created using R (version 3.4.4) with ggplot2 package (version 3.1.0). Sources: Sistema de Vigilância de Agravos de Notificação (SINAN) – Ministry of Health, Brazil, and Instituto Pereira Passos – Rio de Janeiro City Hall, Brazil.

### Dengue, chikungunya, and Zika multivariate clusters

The multivariate scan statistic for multiple data sets detected 17 clusters, of which nine showed dengue, chikungunya, and Zika occurring simultaneously; five showed overlapping dengue and Zika outbreaks; and three showed only outbreaks of Zika (Table 5, Fig 6). The most likely cluster was found in the Downtown region of the city.

**Table 5.**
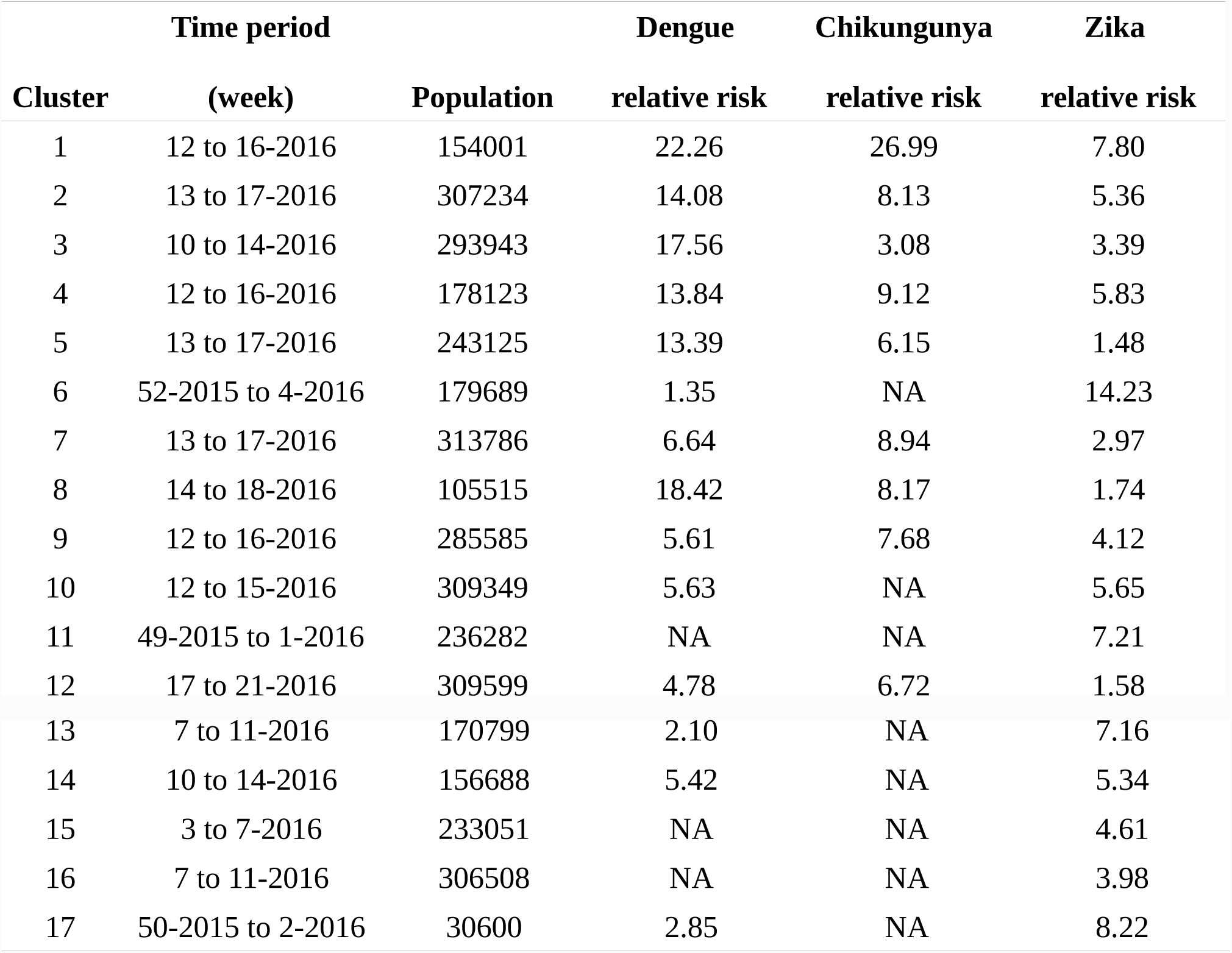
Characteristics of clusters of dengue, chikungunya, and Zika detected using multivariate scan statistic, between epidemiological weeks 30-2015 and 03-2017, Rio de Janeiro city, Brazil. Clusters are ordered according to the maximum likelihood ratio, with 1 being the most likely cluster.

**Fig 6.**
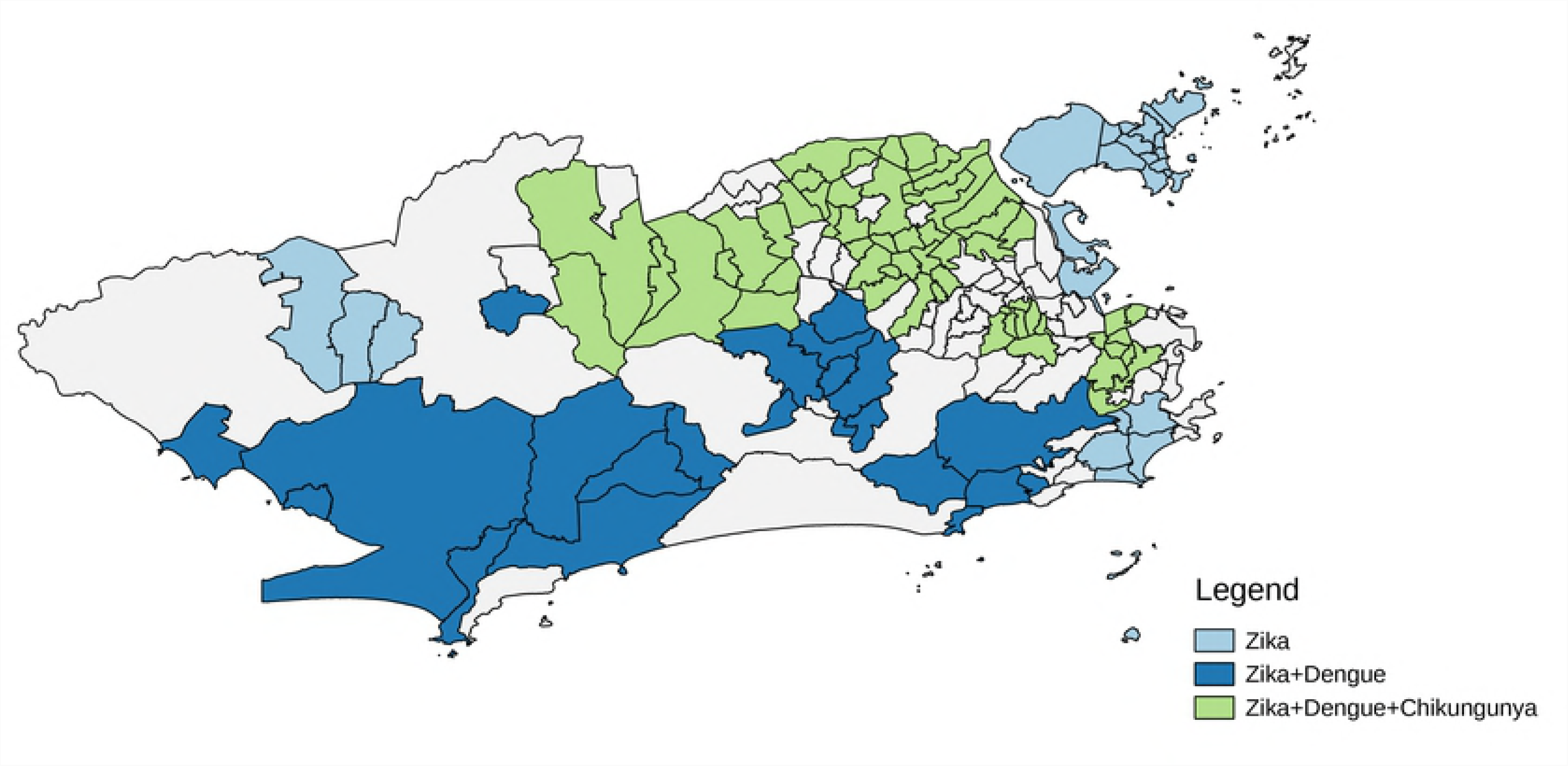
Clusters of dengue, chikungunya, and Zika detected using the multivariate scan statistic, between epidemiological weeks 30-2015 and 03-2017, Rio de Janeiro city, Brazil. Map created using R (version 3.4.4) with ggplot2 package (version 3.1.0). Sources: Sistema de Vigilância de Agravos de Notificação (SINAN) – Ministry of Health, Brazil, and Instituto Pereira Passos – Rio de Janeiro City Hall, Brazil.

Of the 160 neighbourhoods assessed, 57 (35,6%) had clusters for the three diseases coinciding in time and space. Of the nine simultaneous clusters, five were located in the North of the city, three in the West, and one in the Downtown.

## Discussion

This is the first study exploring space-time clustering of dengue, chikungunya, and Zika in an intra-urban region. The data analysed is rare and of great value, as it includes triple epidemics with a large number of cases. Also, this study included the first ever epidemics of chikungunya and Zika in Rio de Janeiro city.

Dengue, chikungunya, and Zika cases were notified across the whole city. The epidemic curves varied slightly in time, with peaks occurring in different weeks. The Zika epidemic curve did not show a clear peak. By stratifying the Zika cases by 10 administrative units of the city (S4 Fig), we hypothesise that the format of the cumulative epidemic curve for the whole city is partially a result of Zika affecting different regions of the city at different times. The number of cases of the three diseases declined after May, coinciding with the end of the rainy and warm season. This reflects the vectors ecology, as *Ae*. *aegypti* and *Ae*. *albopictus* breed in pools of water and temperatures around 25-30°C accelerate the reproductive cycle and increase infectivity and transmissibility [25,26]. In a study in Recife, Northeast Brazil, the simultaneous decrease of Zika and increase of chikungunya cases was also observed. The authors interpreted this as a displacement of Zika caused by chikungunya [27]. For Rio de Janeiro city, this might not be the case, as CHIKV caused only a few cases at beginning of 2016, and only started to rise when Zika cases decreased (the depletion of susceptible hosts). Therefore, we hypothesise that ZIKV circulation inhibited CHIKV, rather than CHIKV introduction displacing ZIKV.

Scan analysis successfully identified clusters of dengue, chikungunya, and Zika. The most likely cluster for each disease occurred in a different part of the city (North, Downtown, and West, respectively). Unlike for dengue and Zika, chikungunya clusters were rarely detected in the West of Rio de Janeiro, probably because the rainy and warm season ended before the disease could reach this region with a sufficient transmission rate to form clusters.

Zika clusters were detected over a longer period of time compared to dengue and chikungunya clusters. We hypothesise that this is a result of the ZIKV advantage in competing for *Ae*. *aegypti* mosquitoes: the *Ae*. *aegypti* has been described as a more efficient vector for ZIKV transmission than for DENV or CHIKV, even when co-infected [28,29]. Not only does *Ae*. *aegypti* transmit ZIKV at a higher rate, but it is also more easily infected by ZIKV compared to DENV and CHIKV. CHIKV, on the other hand, replicates better than ZIKV in *Ae*. *albopictus* cells [28]. While *Ae*. *aegypti* is highly adapted in urban settings, living preferably in domestic and peridomestic areas, *Ae*. *albopictus* prefers to live in areas with more vegetation. However, *Ae*. *albopictus* was recently identified distant from green areas in a densely urbanized complex of favelas in Rio de Janeiro, suggesting this species is adapting to anthropic environments [30]. Further studies are needed to understand the importance of *Ae*. *albopictus* in CHIKV transmission.

A previous study suggested that a Zika epidemic would prevent a subsequent dengue epidemic, as a consequence of cross-immunity [31]. Like DENV, ZIKV is a flavivirus, and the structural similarity between them results in cross-immunity. [32] Whether this cross-immunity leads to antibody-dependent enhancement (ADE, that results in more severe forms of the disease), protection, or neither, is still uncertain [33–35]. In our study, the number of dengue cases increased after the peak of Zika cases. Additionally, some locations with Zika clusters also experienced dengue clusters afterwards. Zika and dengue clusters were spread all over the city. It seems as though herd immunity to dengue did not have a significant impact on the dynamics of Zika or dengue. In the study period, DENV-4 was the most prevalent dengue serotype, followed by DENV-

1. These serotypes were previously responsible for the majority of dengue cases in 2011 (DENV-1) and 2012-2013 (DENV-4). The co-circulation of the 4 dengue serotypes and Zika in the city reinforce the need for active disease surveillance. The consequences of previous DENV exposure to Zika clinical outcomes (and vice-versa) are not clear. By the time the epidemic of congenital Zika syndrome in Brazil was detected, many researchers questioned if it was related to the mother’s anti-DENV antibodies. There is no sufficient evidence to confirm this hypothesis. However, considering the severe consequences of congenital Zika syndrome, disease surveillance using spatio-temporal scan statistics should be considered to identify high risk areas for Zika in a timely manner and to direct preventive measures to the most at risk areas.

Dengue, chikungunya, and Zika clusters detected in Rio de Janeiro do not usually coincided in time and space, contrasting with a study in Mexico that found strong spatio-temporal coherence in the distribution of the three diseases [9]. In addition to virus interactions and competition for the resources for replication inside the vector, behaviour changes may also impact disease dynamics. A rise in the number of cases may promote vector-control activities, which in turn may decrease the number of cases and hinder the establishment of another arbovirus [36]. Also, wealthier areas may have better vector-control interventions, resulting in different spatial distributions.

Neighbourhoods in the North of the city were more likely to have simultaneous clusters of dengue, Zika and chikungunya, highlighted these areas as priority targets for interventions. This is especially important considering co-infections are possible and clinical outcomes are not clear for such cases [37]. As dengue has been endemic in Rio de Janeiro for the last three decades and notification of Zika cases was only established in the municipality in October 2015, it was only possible to detect the first disease cluster for chikungunya and pinpoint its source in the North of the city, highlighting once again the importance of interventions in this area. The North of Rio de Janeiro has already been identified as a hot spot for dengue and as a key region for dengue diffusion. Previous studies also identified Catumbi, a neighbourhood in the Downtown area, as a high-risk location for dengue [38,39]. In our findings, Catumbi comprised the most likely chikungunya cluster, the second most likely cluster for dengue and the third most likely for Zika. Additionally, the clusters in Catumbi coincided in time (most likely cluster in the multivariate scan analysis). Further investigations should be conducted to understand why this neighbourhood in particular is a high-risk location for arboviruses.

The North of the city is marked by a combination of high population density and a lower HDI than the city average [13]. The high population density facilitates the mosquito-human contact and hence the chance of becoming infected. The link between poverty and arbovirus is controversial [40]. Nonetheless, locations with social and economic vulnerability more likely have poorer sanitary conditions and less efficient vector-control interventions, which would facilitate mosquito proliferation. In Rio de Janeiro city, areas in or near favelas were detected as hot spots for dengue [39]. Consistent with our findings, a study conducted in French Guiana indicated that, early in the epidemic, the poorest neighbourhoods would have a greater risk for CHIKV infection [41]. In the first dengue epidemic in a city of São Paulo state, Brazil, authors found a direct relationship between low socio-economic conditions and dengue [42]. We did not observe this relationship for dengue possibly because dengue has already had sustained transmission in the city for decades.

Some limitations affect this study. As our study population included only notified cases (i.e. only patients who sought medical care), asymptomatic cases were not captured. Mild cases usually are poorly captured by SINAN, but considering the disease awareness around Zika, people (especially women) were expected to be more concerned about seeking medical care in case of suspected Zika. As Zika, dengue and chikungunya share some symptoms, the disease awareness may have boosted the notification of mild cases of the three diseases. The similar clinical manifestations of dengue, Zika, and chikungunya also represent a limitation. This limitation is inherent of every study using notified cases, as only a small proportion of cases are laboratory confirmed. However, if misdiagnosis was common, we would not expect to detect differences in time and space of occurrences. In addition, the extensive experience of health care professionals working in Rio de Janeiro, in detecting and diagnosing dengue symptoms, is thought to reduce the probability of misdiagnosis.

A small percentage of cases (8%) that were not georeferenced (and hence, not included in this study) could potentially result in a selection bias. It is possible that cases occurring in favelas, where addresses are sometimes not standardized, have a higher chance of not being georeferenced. Clustering was based on the neighbourhood of residence only, yet infection can happen at other places, such as the workplace. Scan analysis was not designed to understand diseases trajectory but are still helpful to help plan interventions. Also, the method detects circular clusters only, rather than clusters of irregular shapes.

Vector-control strategies have not been effective in abating dengue or in preventing the entry of Zika and chikungunya in Rio de Janeiro. The identification of clusters in space and time allows actions to be intensified in high-risk locations in a timely manner. Special attention should be given to neighbourhoods with high population density and social vulnerability. As vector-control relies on community participation, it is important to enhance community engagement and build trust among all members of the community. People living in neighbourhoods with poor sanitation and a low development index may be less likely to adhere and to maintain prevention activities. Measures to reduce inequity should be accompanied by sustained community engagement [36]. Finally, we suggest the implementation of spatio-temporal scan statistics in the municipal surveillance routine as a tool to optimize prevention strategies.

## Acknowledgements

The authors would like to thank the Municipal Secretariat of Health for providing the data on reported cases, and Dr. Reinaldo Souza dos Santos (Escola Nacional de Saúde Pública Sergio Arouca) and Dr. Valéria Saraceni (Municipal Secretary of Health and Civil Defense, City Hall of Rio de Janeiro) for reviewing and providing helpful feedback.

## Supporting Information

**S1 Fig. Detection of Zika cases clusters according to different temporal and spatial parameters. A) Default parameters. B) Maximum temporal window of 1 week. C) Maximum temporal window of 4 weeks. D) Maximum temporal window of 4 weeks and maximum of 5% of population at risk. E) Maximum temporal window of 4 weeks and maximum of 1% of population at risk.** Maps were created using R (version 3.4.4) with ggplot2 package (version 3.1.0). Sources: Sistema de Vigilância de Agravos de Notificação (SINAN) – Ministry of Health, Brazil, and Instituto Pereira Passos – Rio de Janeiro City Hall, Brazil.

**S2 Fig. Relative risks of clusters of (A) dengue, (B) chikungunya, and (C) Zika, detected between epidemiological weeks 30-2015 and 03-2017 in Rio de Janeiro city, Brazil.** Maps were created using R (version 3.4.4) with ggplot2 package (version 3.1.0). Sources: Sistema de Vigilância de Agravos de Notificação (SINAN) – Ministry of Health, Brazil, and Instituto Pereira Passos – Rio de Janeiro City Hall, Brazil.

**S3 Fig. Week of cluster detection for (A) dengue, (B) chikungunya, and (C) Zika, in Rio de Janeiro city, Brazil.** Maps were created using R (version 3.4.4) with ggplot2 package (version 3.1.0). Sources: Sistema de Vigilância de Agravos de Notificação (SINAN) – Ministry of Health, Brazil, and Instituto Pereira Passos – Rio de Janeiro City Hall, Brazil.

**S4 Fig. Distribution of Zika cases notifications by week and administrative units (programmatic area – AP) of Rio de Janeiro city city.** Source: Sistema de Vigilância de Agravos de Notificação (SINAN) – Ministry of Health, Brazil.

